# Persistence of a *Wolbachia*-driven sex ratio bias in an island population of *Eurema* butterflies

**DOI:** 10.1101/2020.03.24.005017

**Authors:** Daisuke Kageyama, Satoko Narita, Tatsuro Konagaya, Mai N. Miyata, Jun Abe, Wataru Mitsuhashi, Masashi Nomura

## Abstract

It is generally believed that when maternally inherited sex ratio distorters become predominant, either the host population goes extinct or nuclear suppressors evolve in the host. Here, we show an empirical case where all-female-producing *Wolbachia* is likely to be stably maintained at a high frequency. On an island population of the butterfly *Eurema mandarina*, a *Wolbachia* strain *w*Fem, which makes female hosts produce all-female offspring without sibling lethality (female drive), is highly prevalent. We found that, with some fluctuations, *w*Fem appeared to be stably maintained for at least 12 years at a high frequency, resulting in the existence of an abnormally high number of virgin females. Interestingly, comparison between sex ratios of captive individuals and sex ratios deduced from *w*Fem frequencies suggested a plastic behavioral change of males and females in response to the shift of sex ratios. *w*Fem presence does not affect brood size but has a slightly negative effect on body size. Stable coexistence of *w*Fem-positive and -negative females in the population may be explained via mate choice by males, which keeps *w*Fem in check. Taken together, this butterfly population is an attractive model for future studies on the population dynamics of sex ratios and mating behavior.

## Introduction

Arthropods are often influenced by selfish microbes, such as *Wolbachia*, which are maternally transmitted through the cytoplasm and can skew the sex ratio toward females (Hurst and Jiggins 2000; Werren et al. 2008; Kageyama et al. 2012). It is predicted that the prevalence of cytoplasmic sex ratio distorters can lead to the extinction of the host population (Hatcher et al. 1999), unless suppressors evolve in the host. To date, the existence of host nuclear suppressors against cytoplasmic sex ratio distorters has been described in various arthropod species (Jaenike 2007; Majerus and Majerus 2010), and the rapid spread of suppressors in natural populations has been documented in two species—i.e., the butterfly *Hypolimnas bolina* (Lepidoptera; Nymphalidae) (Mitsuhashi et al. 2004; Hornett et al. 2006; Charlat et al. 2007a; Mitsuhashi et al. 2011) and the lacewing *Mallada desjardinsi* (Neuroptera; Chrysopidae) (Hayashi et al. 2016, 2018)—which suggests that an arms race concerning the sex ratio between the cytoplasmic and nuclear genomes may be quite common in insects.

Here, we examined the sex ratio dynamics in an island population (Tanegashima Island, Japan) of the butterfly *Eurema mandarina* (Lepidoptera: Pieridae), where all-female-producing females and normal females coexist. In *E. mandarina*, cytoplasmic incompatibility-inducing *Wolbachia* (*w*CI), which does not influence the sex ratio, is fixed in most of the populations in Japan, including the Tanegashima Island population (Hiroki et al. 2005; Narita et al. 2006). By contrast, all-female-producing females, which are infected with another strain of *Wolbachia*, *w*Fem, besides the *w*CI strain (Hiroki et al. 2002, 2004), are prevalent in the Tanegashima Island population (Narita et al. 2007a, b; Miyata et al. 2017).

In *E. mandarina*, females and males singly infected with *w*CI (referred to as C females and C males) have the WZ karyotype and the ZZ karyotype, respectively, whereas females doubly infected with *w*CI and *w*Fem (referred to as CF females) have the Z0 karyotype (Kern et al. 2015; Kageyama et al. 2017). By mating with C males, CF females produce only Z0 embryos; all of which develop into females. Unlike the male killing effect induced by many *Wolbachia* strains, *w*Fem does not induce male-specific lethality. From a population genetics perspective, this phenomenon can be regarded as female drive (Burt and Trivers 2006), and thus, CF females will increase their frequency in the population if the relative fitness of CF females is higher than that of 50% of C females.

If CF females continue to increase their frequency in the population, the population could become extinct because the dwindling number of males would become too small to maintain the population. Alternatively, nuclear suppressors against female drive, if they arise, may rapidly increase in the population. With these possible scenarios in mind, we monitored the island population of *E. mandarina* for 12 years.

## Materials and Methods

### COLLECTION OF BUTTERFLIES

On Tanegashima Island (Kagoshima Prefecture, Japan), we patrolled the pavement and made efforts to collect all *E. mandarina* adults that we encountered. Males were stored at −30°C until DNA extraction began. After allowing females to lay eggs in the laboratory for other experiments (Narita et al. 2007a; Kageyama et al. 2017), bursa copulatrixes were dissected out from the female specimens; the remains were stored at −30°C until DNA extraction began.

### DISSECTION OF SPERMATOPHORES

As described in Konagaya and Watanabe (2015), the bursa copulatrixes were carefully opened under a dissecting microscope, and the number of spermatophores that each contained was recorded.

### MEASUREMENT OF WING SIZE

For adult females, the distance between the tip of the forewing and the base of the forewing jointed with the thorax was measured.

### DNA EXTRACTION AND PCR

DNA was extracted from either specimen legs or thoraxes using a DNeasy Blood & Tissue Kit (Qiagen, Tokyo, Japan). *Wolbachia* infections of either the *w*CI or *w*Fem strain were identified via specific PCR detection, which targeted the *wsp* gene (Hiroki et al. 2004; Narita et al. 2007b; Kageyama et al. 2008). Specifically, *w*CI was detected using the *Wolbachia*-specific forward primer wsp81F (5’-TGGTCCAATAAGTGATGAAGAAAC-3’) (Braig et al. 1998) and the *w*CI-specific reverse primer WHecFem1 (5’-ACTAACGTTTTTGTTTAG-3’) (Hiroki et al. 2004), which amplify a 232 bp DNA fragment. The *w*Fem strain was detected using the *w*Fem-specific forward primer WHecFem2 (5’-TTACTCACAATTGGCTAAAGAT-3’) (Hiroki et al. 2004) and the *Wolbachia*-specific reverse primer wsp691R (5’-AAAAATTAAACGCTACTCCA-3’) (Braig et al. 1998), which amplify a 398 bp DNA fragment.

### STATISTICAL ANALYSES

Analyses of binary data (i.e., virgin or mated females) were conducted using generalized linear mixed models (GLMM) with a binomial error distribution. Analyses of mating frequencies (i.e., number of spermatophores per female) were conducted using GLMM with a Poisson error distribution. The random effects of different visits were included in the models. The GLMM were fitted using Laplace approximation. All statistical analyses were performed using the function *glmer* of the program package lme4 using R version 3.4.4 (R Core Team 2017).

## Results and Discussion

### CF FEMALES CONSTANTLY PRODUCE ALL OR NEARLY ALL-FEMALE OFFSPRING

As was partially shown previously (Narita et al. 2007a,b; Miyata et al. 2017), all but one of the 36 CF females collected on Tanegashima Island produced only daughters (1024 females and no males for 35 broods, and seven females and one male for one brood; Table S1), whereas all 17 C females produced sons and daughters (211 females and 220 males for the recorded 13 broods; Table S1). These results confirm that suppressors against female drive are absent or very low in frequency, if present. It is also likely that there are no sex ratio distorters other than *w*Fem in the population.

Because the all-female trait of CF matrilines is not due to male killing, the number of daughters produced by CF females was nearly twice as much as that produced by C females (Kern et al. 2015; Kageyama et al. 2017) (Table S1). Vertical transmission efficiencies of *w*CI and *w*Fem would be high because both *w*CI and *w*Fem were detected by PCR from all 40 daughters from each of the three all-female broods produced by females collected in 2007 (*n* = 120) as well as from 12, 14, 11, and 11 daughters from the four all-female broods produced by females collected in 2015 (*n* = 48). Taken together, CF females would be expected to increase in the population until the population became extinct unless some counterforces existed.

### FEMALE-BIASED POPULATION SEX RATIO IS ASSOCIATED WITH A HIGH FREQUENCY OF CF FEMALES

In total, 668 adults were captured on Tanegashima Island, Japan, from 2005 to 2017 (through 13 visits), which included 523 females and 145 males. Sex ratios varied among the 13 visits (37.9%–100% female; median = 86.5%; Table S2). Out of 495 females diagnosed with *Wolbachia* infection, 495 (100%) tested positive for the *w*CI strain and 415 (83.8%) were also positive for the *w*Fem strain (i.e., 80 C females and 415 CF females; Table S2).

The proportion of CF females among females (PCF) varied among the 13 visits (54.2%–92.5%; median = 84.4%; Table S2). By assuming that (1) equal numbers of C females and males existed in the population and (2) behavioral activity did not differ between C and CF females, the proportion of females (P_female_) can be deduced from P_CF_ as

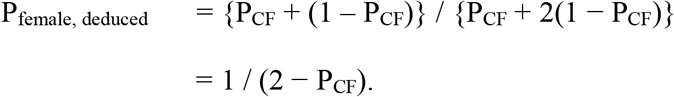

For the 13 visits, these deduced sex ratios (P_female, deduced_) were compared with the observed sex ratios (i.e., P_female, observed_; proportion of females among captive individuals) (Figure 1A). These values fit well, except for four visits with the least biased sex ratios (the 3^rd^, 6^th^, 10^th^, and 11^th^ visits; values were only significantly different on the 10^th^ and 11^th^ visits) and three visits with extremely female-biased sex ratios (the 1^st^, 2^nd^, and 13^th^ visits; values were only significantly different on the 2^nd^ visit). We speculate that these discrepancies are due to a temporary change in male behavior in response to fluctuation of the sex ratio. When males are not extremely rare, males are more conspicuous and easily recognized by investigators than females because males actively search for females, one after another, despite frequent rejections. This is also a typical situation in *E. mandarina* populations without CF females, where the actual sex ratio is 1:1 (observed male to female ratios were 1.5–17 in seven populations; median, 3.22; Figure 1B). A male-biased sex ratio of captive individuals has also been reported in other butterfly species (Matsumoto 1984; Kitamura 1999; Kobayashi and Inaizumi 2003). However, males become less conspicuous under extremely female-biased conditions with a high proportion of virgin females because males would have high rates of mating success. In such extreme conditions, males may become less conspicuous than females; in such a case, females may actively search for males. During a visit in 2005, one team member described that females were actively flying, possibly searching for males, and that such behavior is unusual in other *E. mandarina* populations.

**Figure 1.**
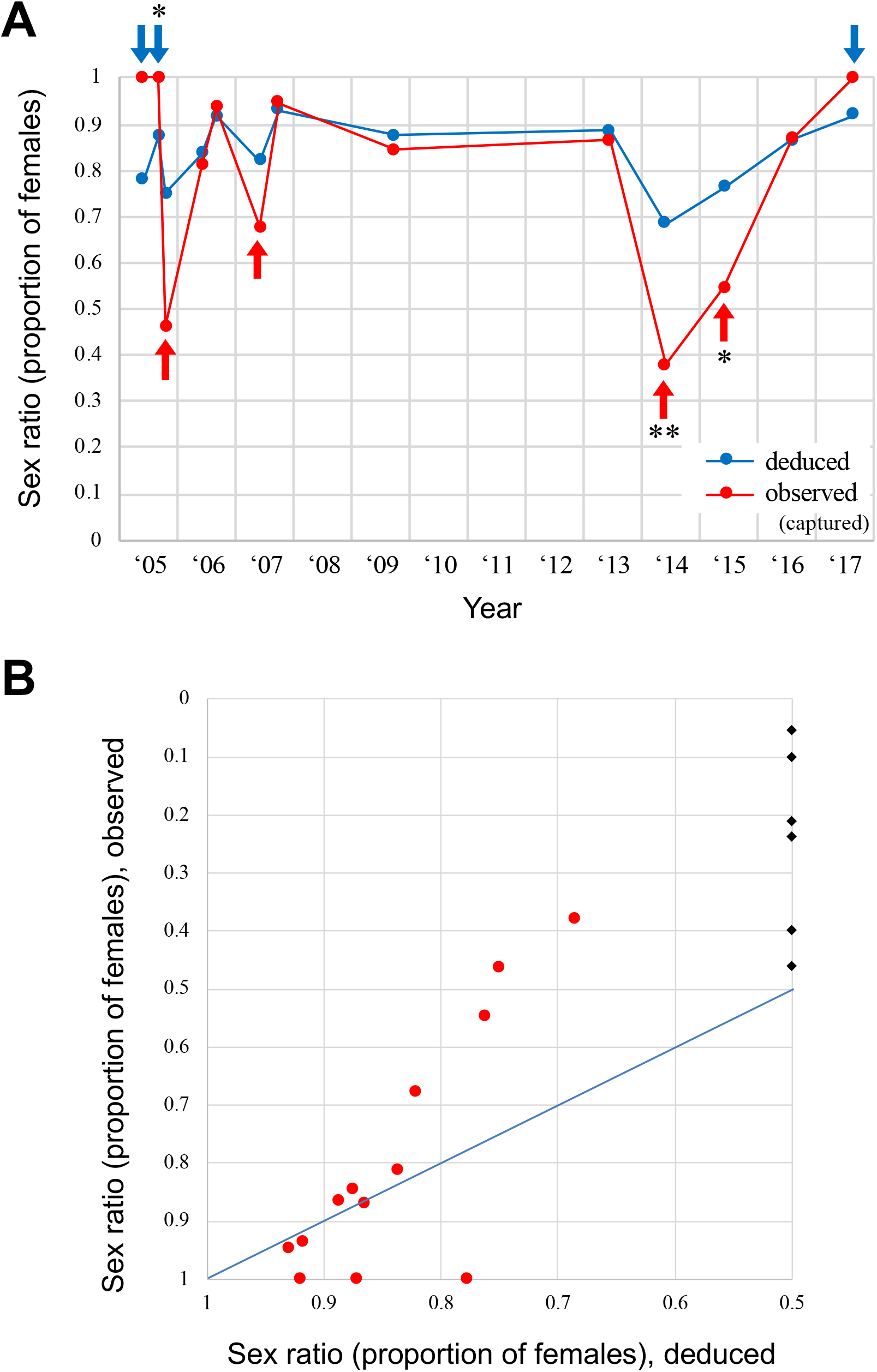
Sex ratios of *E. mandarina* on Tanegashima Island. (A) Fluctuation of observed sex ratios (i.e., the proportions of females among captive individuals; given in red) and deduced sex ratios (i.e., the proportions of females deduced from infection frequencies of *w*Fem; given in blue) examined in 13 visits from 2005 to 2017. Four visits, where males were overrepresented in the observed sex ratios, are indicated by red arrows. Three visits, where males were underrepresented in the observed sex ratios, are indicated by blue arrows. Asterisks indicate a significant difference between observed and deduced sex ratios by the chi-squared test, which compared the actual numbers of captive males and females versus the numbers of *Wolbachia*-diagnosed females (C and CF females) and the virtual number of males (by assuming that it is equal to the number of C females) (\**p* < 0.05; \*\**p* < 0.01 by the chi-squared test). (B) Correlation between observed sex ratios and deduced sex ratios. Red circles: data from 13 visits to Tanegashima Island. Black rhombi: data from six other populations without the *w*Fem strain. The blue line shows where the observed and deduced sex ratios are equal.

This behavioral change would thus be plastic; indeed, turnovers of observed and deduced sex ratios were observed multiple times (Figure 1A). However, female-biased conditions, if enduring for long enough, may select for genetic traits such as those of males with less active mating behavior or females with more active mating behavior. In the butterfly *Acrea encedon* (Lepidoptera: Nymphalidae), high prevalence of male-killing *Wolbachia* is considered to have resulted in sex role reversal concerning mating behavior (Jiggins et al. 2000). Some slight genetic changes in mating behavior may exist in the Tanegashima Island population compared with other populations without *w*Fem.

It is unclear when CF females first invaded Tanegashima Island. According to the butterfly inventory, with records that date back to the 1960s, 16 females and 24 males were observed between 1960 and 1978 and 17 females and 16 males were observed between 2002 and 2007 (no significant difference between the two periods was detected by the chi-squared test; Table S3). Importantly, an author noted in their graduate thesis that one of the samples collected from Tanegashima Island in 1998 was PCR-positive for *w*Fem. Although we have no idea whether *w*Fem was present in 1960–1978, it is likely that *w*Fem was already present and spread in 2002–2004, before the start of the present study.

### SCARCITY OF MALES LEADS TO A HIGH PROPORTION OF VIRGIN FEMALES

The proportion of virgin females is abnormally high on Tanegashima Island (12.2%–82.6%; median = 41.5%) compared with that of other *E. mandarina* populations that lack CF females (0%–4.76%; median = 0%) (Table 1). Overall, the proportion of virgin females was significantly correlated with the sex ratio (*P* = 0.0124 by GLMM), but they were discrete depending on the season (Figure 2A). Significant correlations between the sex ratio and virginity were observed for females collected in June (*P* = 0.00554 by GLMM) and for females collected in September (*P* = 0.0102 by GLMM) (Figure 2A). Similarly, the number of spermatophores per female was significantly correlated with the sex ratio (*P* = 0.000724), but they were discrete depending on the season (Figure 2B). Significant correlations between the sex ratio and number of spermatophores were observed for females collected in June (*P* = 0.0379 by GLMM) and for females collected in September (*P* = 0.0179 by GLMM) (Figure 2B). Taken together, it is likely that the *Wolbachia*-induced scarcity of males resulted in a low mating frequency on Tanegashima Island. The higher mating tendency in late summer (September) compared with early summer (June) could be explained by higher male activity because of the elevated temperature. The larger population size in late summer compared with early summer may also increase the chance of encounters between males and females regardless of the sex ratio.

**Table 1.** Number of spermatophores possessed by wild-caught *E. mandarina* females

|  | Proportion of females | No. of females with *n* spermatophore(s) |  |  |  |  |  |  | Proportion of virgin females | Mean no. of spermatophores per female |
| --- | --- | --- | --- | --- | --- | --- | --- | --- | --- | --- |
|  |  | *n* = 0 (virgin) | *n* = 1 | *n* = 2 | *n* = 3 | *n* = 4 | *n* = 5 | *n* = 6 |  |  |
| **(a) Tanegashima population** |  |  |  |  |  |  |  |  |  |  |
| Tanegashima (Sep, 2005) | 1.000 (42) | 25 | 9 | 0 | 0 | 0 | 0 | 0 | 0.735 | 0.265 |
| Tanegashima (Jun, 2006) | 0.811 (53) | 17 | 21 | 2 | 1 | 0 | 0 | 0 | 0.415 | 0.683 |
| Tanegashima (Sep, 2006) | 0.936 (109) | 32 | 48 | 18 | 0 | 0 | 0 | 0 | 0.327 | 0.857 |
| Tanegashima (Jun, 2007) | 0.676 (34) | 3 | 14 | 3 | 0 | 0 | 0 | 0 | 0.150 | 1.000 |
| Tanegashima (Sep, 2007) | 0.947 (57) | 20 | 25 | 6 | 2 | 0 | 0 | 0 | 0.377 | 0.811 |
| Tanegashima (Sep, 2009) | 0.845 (58) | 6 | 33 | 7 | 3 | 0 | 0 | 0 | 0.122 | 1.143 |
| Tanegashima (Jun, 2013) | 0.865 (74) | 35 | 25 | 2 | 0 | 0 | 1 | 0 | 0.556 | 0.540 |
| Tanegashima (Jul, 2016) | 0.868 (38) | 16 | 16 | 0 | 1 | 0 | 0 | 0 | 0.485 | 0.576 |
| Tanegashima (Aug, 2017) | 1.000 (24) | 19 | 3 | 0 | 1 | 0 | 0 | 0 | 0.826 | 0.261 |
|  |  |  |  |  |  |  |  |  | 0.415 (median) | 0.683 (median) |
| **(b) Other populations** |  |  |  |  |  |  |  |  |  |  |
| Zao, Miyagi Pref. (Sep, 2006) | 0.400 (20) | 0 | 4 | 1 | 0 | 2 | 1 | 0 | 0.000 | 2.375 |
| Tochigi, Tochigi Pref. (Sep, 2009) | 0.262 (61) | 0 | 1 | 3 | 3 | 2 | 0 | 1 | 0.000 | 3.000 |
| Shodoshima, Kagawa Pref. (Aug, 2011) | 0.237 (135) | 0 | 9 | 8 | 6 | 0 | 0 | 0 | 0.000 | 1.870 |
| Hachijojima (Sep, 2011) | 0.460 (139) | 3 | 29 | 18 | 10 | 2 | 1 | 0 | 0.048 | 1.714 |
|  |  |  |  |  |  |  |  |  | 0.000 (median) | 2.122 (median) |

**Figure 2.**
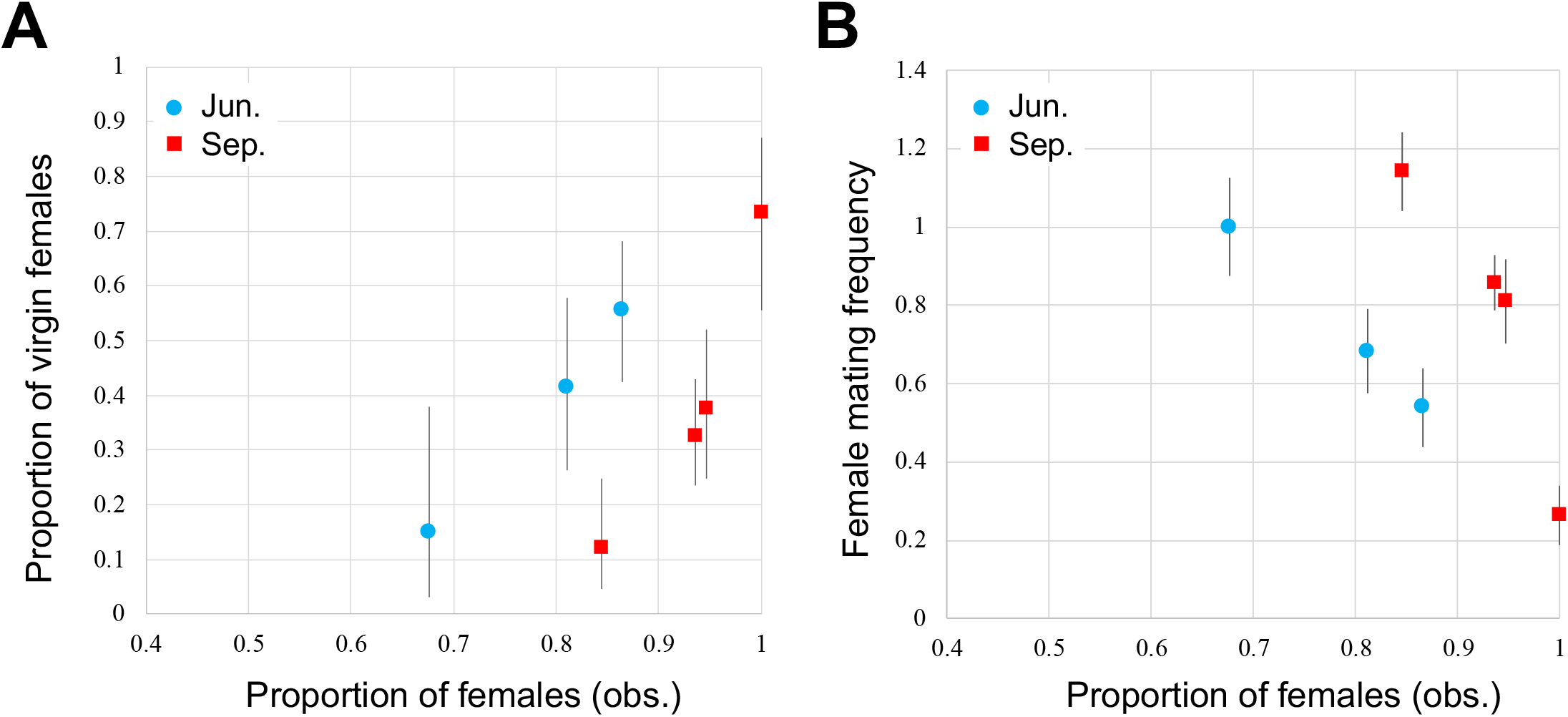
Correlation between sex ratios and mating tendency of *E. mandarina* on Tanegashima Island. (A) Proportions of virgin females plotted against observed sex ratios (proportions of females). The error bars indicate 95% confidence intervals. (B) Female mating frequencies (i.e., numbers of spermatophores per female) plotted against the observed sex ratio. The error bars indicate the standard error of the mean. Plots for females collected in June are represented by blue circles, and those collected in September are represented by red squares.

In *H. bolina*, *Wolbachia*-induced female bias leads to an increase in female mating frequency, up to a point where male mating capacity becomes a limiting factor (Charlat et al. 2007b). The increased female mating frequency is considered to be a facultative response to depleted male mating resources in female-biased populations. In *E. mandarina*, alternatively, female mating frequency did not seem to increase in response to the shift of the sex ratio toward females (Figure 2B), which may indicate that this could be explained by the difference in male mating capacity between the two species.

### COSTS OF *w*FEM INFECTION

Although we could not detect a significant difference in the number of offspring between wild-caught C and CF females (*P* = 0.748 by GLM; Table S1), our data concerning wing size suggest that the *w*Fem strain has some physiological costs. Although female wing size tended to be larger in June than in September, which may be associated with prolonged hormonal events at lower temperatures (Davidowitz et al. 2004), the wing size of CF females was significantly smaller than that of C females (*P* = 0.042 by GLMM; Table S4). Narita et al. (2007b) also found that the offspring of CF females required longer development times than the offspring of C females. Although more detailed experiments will be necessary in future work, it is likely that *w*Fem imposes some fitness costs to *E. mandarina*.

### MAINTENANCE OF *w*FEM IN THE POPULATION

We do not believe that *w*Fem reduces fitness; otherwise, the invasion of CF females in the population could not be explained in the first place. The stable maintenance of female polymorphism (i.e., the coexistence of C and CF females in the population) for 12 years (more than 48 generations) (Kato 1986) could have been achieved by suppressing the increase of CF females, possibly via mate choice by males (Bonduriansky 2001) in favor of C females; this is plausible because C females are larger than CF females. Males choose larger females in some butterfly species (Marshall 1982; Rutowski et al. 1983; Van Dongen et al. 1998), including those in the genus *Eurema* (Kemp 2008). One scenario, the so-called negative frequency-dependent selection, assumes genetic polymorphism in males regarding mating behavior (i.e., the existence of choosy and non-choosy males) and fitness costs in choosy males. When the sex ratio is female-biased, producing sons is more advantageous than producing daughters (Fisher 1930; Hamilton 1967). Hence, choosy males that favor C females will be selected for because they produce a larger number of sons than non-choosy males. Upon an increase in relative frequency of C females, the sex ratio becomes less biased, which is a scenario where selection for choosy males becomes relaxed. Then, choosy males will decrease their frequency because of the fitness cost, which allows CF females to increase.

Another scenario that we consider more plausible is one that does not assume genetic polymorphism. Males may change their mating behavior in response to the shift of the population sex ratio. In a situation where males are not very rare, males may indiscriminately mate with both types of females on a first-come basis. This situation will increase the relative frequency of CF females, and males will become increasingly rare. At some point, males may often encounter multiple females simultaneously, and because males tend to choose larger females (i.e., C females) (Kemp 2008), the relative frequency of C females, and males, will increase.

However, our study shows that the proportion of virgin females and the numbers of spermatophores were not significantly different between CF and C females (*P* = 0.599 for the proportion of virgins; *P* = 0.658 for the numbers of spermatophores by GLMM) (Table S5). If mating discrimination only occurs when the sex ratio is extremely biased toward females, the extremely small number of C females would limit the statistical power to detect the potential existence of mate choice. More thorough sampling of females and behavioral experiments may reveal the abovementioned possibilities in our future work.

Although less likely, the fixation of CF females might be hindered by the periodical and constant immigration of C females onto Tanegashima Island from other populations. Furthermore, the metapopulation structure may explain the coexistence of C and CF females in Tanegashima Island. Thus, it would be of interest to examine *E. mandarina* genetic diversity in the Tanegashima Island population as well as other populations without CF females to determine the validity of these distinct hypotheses. If migration is very rare, this population may suffer from substantial inbreeding resulting from the sex ratio bias (i.e., a small effective population size).

## Conclusions

Despite our initial expectation of either population decline, extinction, or the spread of suppressors, the polymorphism (i.e., the coexistence of C and CF females) appears to have been maintained in the population for at least the last 12 years (>48 generations) without any evidence of suppressors. Our data may be explained by the cyclic dynamics of C and CF females in the population, because of either negative frequency-dependent selection for male genetic polymorphisms or a plastic behavioral change in response to the shift of sex ratios. Longer-term effects of the sex ratio bias, such as genetic changes in mating behavior and substantial inbreeding due to the small effective population size, may also exist.

## LITERATURE CITED

Bonduriansky, R. 2001. The evolution of male mate choice in insects: a synthesis of ideas and evidence. Biol. Rev. 76:305–339.

Braig, H. R., W. Zhou, S. L. Dobson, and S. L. O’Neill. 1998. Cloning and characterization of a gene encoding the major surface protein of the bacterial endosymbiont *Wolbachia pipientis*. J. Bacteriol. 180:2373–2378.

Burt, A., and R. Trivers. 2006. Genes in Conflict - The Biology of Selfish Genetic Elements. Harvard University Press, London, UK.

Charlat, S., E. A. Hornett, J. H. Fullard, N. Davies, G. K. Roderick, N. Wedell, and G. D. D. Hurst. 2007a. Extraordinary flux in sex ratio. Science 317:214–214.

Charlat, S., M. Reuter, E. A. Dyson, E. A. Hornett, A. Duplouy, N. Davies, G. K. Roderick, N. Wedell, and G. D. D. Hurst. 2007b. Male-killing bacteria trigger a cycle of increasing male fatigue and female promiscuity. Curr. Biol. 17:273–277.

Davidowitz, G., L. J. D’Amico, and H. F. Nijhout. 2004. The effects of environmental variation on a mechanism that controls insect body size. Evol. Ecol. Res. 6:49–62.

Fisher, R. A. 1930. The Genetical Theory of Natural Selection. Clarendon Press, Oxford, UK.

Hamilton, W. D. 1967. Extraordinary sex ratios. Science 156:477–488.

Hatcher, M. J., D. E. Taneyhill, A. M. Dunn, and C. Tofts. 1999. Population dynamics under parasitic sex ratio distortion. Theor. Pop. Biol. 56:11–28.

Hayashi, M., M. Nomura, and D. Kageyama. 2018. Rapid comeback of males: evolution of male-killer suppression in a green lacewing population. Proc. R. Soc. B 285:20180369.

Hayashi, M., M. Watanabe, F. Yukuhiro, M. Nomura, and D. Kageyama. 2016. A nightmare for males? A Maternally transmitted male-killing bacterium and strong female bias in a green lacewing population. PLoS One 11:e0155794.

Hiroki, M., Y. Ishii, and Y. Kato. 2005. Variation in the prevalence of cytoplasmic incompatibility-inducing *Wolbachia* in the butterfly *Eurema hecabe* across the Japanese archipelago. Evol. Ecol. Res. 7:931–942.

Hiroki, M., Y. Kato, T. Kamito, and K. Miura. 2002. Feminization of genetic males by a symbiotic bacterium in a butterfly, *Eurema hecabe* (Lepidoptera: Pieridae). Naturwissenschaften 89:167–170.

Hiroki, M., Y. Tagami, K. Miura, and Y. Kato. 2004. Multiple infection with *Wolbachia* inducing different reproductive manipulations in the butterfly *Eurema hecabe*. Proc. R. Soc. B 271:1751–1755.

Hornett, E. A., S. Charlat, A. M. R. Duplouy, N. Davies, G. K. Roderick, N. Wedell, and G. D. D. Hurst. 2006. Evolution of male-killer suppression in a natural population. PLoS Biol. 4:e283.

Hurst, G. D., and F. M. Jiggins. 2000. Male-killing bacteria in insects: mechanisms, incidence, and implications. Emerg. Infect. Dis. 6:329–336.

Jaenike, J. 2007. Spontaneous Emergence of a New Wolbachia Phenotype. Evolution 61:2244–2252.

Jiggins, F. M., G. D. Hurst, and M. E. Majerus. 2000. Sex-ratio-distorting Wolbachia causes sex-role reversal in its butterfly host. Proc. R. Soc. B 267:69–73.

Kageyama, D., S. Narita, and H. Noda. 2008. Transfection of feminizing *Wolbachia* endosymbionts of the butterfly, *Eurema hecabe*, into the cell culture and various immature stages of the silkmoth, *Bombyx mori*. Microb. Ecol. 56:733.

Kageyama, D., S. Narita, and M. Watanabe. 2012. Insect sex determination manipulated by their endosymbionts: incidences, mechanisms and implications. Insects 3:161–199.

Kageyama, D., M. Ohno, T. Sasaki, A. Yoshido, T. Konagaya, A. Jouraku, S. Kuwazaki, H. Kanamori, Y. Katayose, S. Narita, M. Miyata, M. Riegler, and K. Sahara. 2017. Feminizing *Wolbachia* endosymbiont disrupts maternal sex chromosome inheritance in a butterfly species. Evol. Lett. 1:232–244.

Kato, Y. 1986. The prediapause copulation and its significance in the butterfly *Eurema hecabe*. J. Ethol. 4:81–90.

Kemp, D. J. 2008. Female mating biases for bright ultraviolet iridescence in the butterfly Eurema hecabe (Pieridae). Behav Ecol 19:1–8. Oxford Academic.

Kern, P., J. M. Cook, D. Kageyama, and M. Riegler. 2015. Double trouble: combined action of meiotic drive and Wolbachia feminization in Eurema butterflies. Biol. Lett. 11:20150095.

Kitamura, M. 1999. Butterflies from the southwest side slope of Mt. Banahaw, Mid-south Luzon, Philippines (1) Nymphalidae. Butterflies 22:9–24 (in Japanese with English summary).

Kobayashi, T., and M. Inaizumi. 2003. The sex ratios of the wild adult populations in *Sasakia charonda* (Nymphalidae). Lepid. Sci. 54:156–162.

Konagaya, T., and M. Watanabe. 2015. Adaptive significance of the mating of autumn-morph females with non-overwintering summer-morph males in the Japanese Common Grass Yellow, *Eurema mandarina* (Lepidoptera: Pieridae). Appl. Entomol. Zool. 50:41–47.

Majerus, T. M. O., and M. E. N. Majerus. 2010. Intergenomic Arms Races: Detection of a Nuclear Rescue Gene of Male-Killing in a Ladybird. PLoS Pathog. 6:e1000987.

Marshall, L. 1982. Male Courtship Persistence in Colias philodice and C. eurytheme (Lepidoptera: Pieridae). J. Kansas Entomol. Soc. 55:729–736.

Matsumoto, K. 1984. Population dynamics of *Luehdorfia japonica* Leech (Lepidoptera: Papilionidae). Res. Pop. Ecol. 26:1–12.

Mitsuhashi, W., H. Fukuda, K. Nicho, and R. Murakami. 2004. Male-killing *Wolbachia* in the butterfly *Hypolimnas bolina*. Entomol. Exp. Appl. 112:57–64.

Mitsuhashi, W., H. Ikeda, and M. Muraji. 2011. Fifty-year trend towards suppression of *Wolbachia*-induced male-killing by its butterfly host, *Hypolimnas bolina*. J. Insect Sci. 11:92.

Miyata, M., T. Konagaya, K. Yukuhiro, M. Nomura, and D. Kageyama. 2017. *Wolbachia*-induced meiotic drive and feminization is associated with an independent occurrence of selective mitochondrial sweep in a butterfly. Biol. Lett. 13:20170153.

Narita, S., D. Kageyama, M. Nomura, and T. Fukatsu. 2007a. Unexpected mechanism of symbiont-induced reversal of insect sex: feminizing *Wolbachia* continuously acts on the butterfly *Eurema hecabe* during larval development. Appl. Environ. Microbiol. 73:4332–4341.

Narita, S., M. Nomura, and D. Kageyama. 2007b. Naturally occurring single and double infection with *Wolbachia* strains in the butterfly *Eurema hecabe*: transmission efficiencies and population density dynamics of each *Wolbachia* strain. FEMS Microbiol. Ecol. 61:235–245.

Narita, S., M. Nomura, Y. Kato, and T. Fukatsu. 2006. Genetic structure of sibling butterfly species affected by *Wolbachia* infection sweep: evolutionary and biogeographical implications. Mol. Ecol. 15:1095–1108.

R Core Team. 2017. R: A language and environment for statistical computing. R Foundation for Statistical Computing, Vienna, Austria.

Rutowski, R. L., M. Newton, and J. Schaefer. 1983. Interspecific Variation in the Size of the Nutrient Investment Made by Male Butterflies During Copulation. Evolution 37:708–713.

Van Dongen, S., E. Matthysen, E. Sprengers, and A. A. Dhondt. 1998. Mate Selection by Male Winter Moths Operophtera brumata (Lepidoptera, Geometridae): Adaptive Male Choice or Female Control? Behaviour 135:29–42.

Werren, J. H., L. Baldo, and M. E. Clark. 2008. *Wolbachia:* master manipulators of invertebrate biology. Nat. Rev. Microbiol. 6:741–751.

