## Supplementary material for "Persistence of a *Wolbachia*-driven sex ratio bias in an island population of *Eurema* butterflies": Table S1

Infection status of *Wolbachia* of wild-caught females and their offspring sex ratio in Tanegashima Island population of *E. mandarina* .

| Wild-caught females | Offspring emerged as adults | | |
|---|---|---|---|
| | Female | Male | Total |
| **(A) Doubly infected (CF)** | | | |
| 2006-Jun-Y01[a] | 37 | 0 | 37 |
| 2006-Jun-O01[a] | 7 | 1 | 8 |
| 2006-Jun-D01[a] | 46 | 0 | 46 |
| 2006-Jun-h01[a] | 14 | 0 | 14 |
| 2006-Jun-Z06[a] | 7 | 0 | 7 |
| 2007-Jun-A02[a] | 16 | 0 | 16 |
| 2007-Jun-M11 | 113 | 0 | 113 |
| 2007-Jun-U03 | 98 | 0 | 98 |
| 2007-Jun-X30 | 83 | 0 | 83 |
| 2007-Jun-Z78 | 64 | 0 | 64 |
| 2007-Jun-Z79 | 77 | 0 | 77 |
| 2007-Jun-Z80 | 11 | 0 | 11 |
| 2013-13a | 19 | 0 | 19 |
| 2013-Q18 | 15 | 0 | 15 |
| 2013-Q39 | 4 | 0 | 4 |
| 2013-Q42 | 6 | 0 | 6 |
| 2014-519Q1 | 17 | 0 | 17 |
| 2014-519Q2 | 11 | 0 | 11 |
| 2014-519Q7 | 15 | 0 | 15 |
| 2014-519Q11 | 10 | 0 | 10 |
| 2014-519Q12 | 7 | 0 | 7 |
| 2014-520Q1 | 13 | 0 | 13 |
| 2014-521Q1 | 10 | 0 | 10 |
| 2014-521Q4 | 4 | 0 | 4 |
| 2014-521HN1 | 6 | 0 | 6 |
| 2014-521HN2 | 19 | 0 | 19 |
| 2014-522Q1 | 17 | 0 | 17 |
| 2015-F-01[b] | 34 | 0 | 34 |
| 2015-F-07[b] | 25 | 0 | 25 |
| 2015-F-08[b] | 42 | 0 | 42 |
| 2015-F-10[b] | 44 | 0 | 44 |
| 2015-F-19[b] | 81 | 0 | 81 |
| 2015-F-20[b] | 35 | 0 | 35 |
| 2015-F-21[b] | 16 | 0 | 16 |
| 2015-F-35[b] | 4 | 0 | 4 |
| 2016-F19 | 4 | 0 | 4 |
| Total (*n* = 36) | 1031 | 1 | 1032 |
| Mean | 28.639 | 0.028 | 28.667 |
| S.D. | 29.211 | 0.167 | 29.190 |
| S.E. | 4.868 | 0.028 | 4.865 |
| Median | 16 | 0 | **16** |
| **(B) Singly infected (C)** | | | |
| 2013-shimonishi-1 | 14 | 20 | 34 |
| 2013-ishido-1 | 2 | 4 | 6 |
| 2014-519Q8 | 4 | 10 | 14 |
| 2014-519Q10 | 9 | 6 | 15 |
| 2014-519Q13 | 7 | 12 | 19 |
| 2014-519Q15 | 7 | 10 | 17 |
| 2014-521minami-1 | 7 | 7 | 14 |
| 2015-F-05 | 58 | 40 | 98 |
| 2015-F-06 | 8 | 7 | 15 |
| 2015-F-27 | 31 | 52 | 83 |
| 2015-F-28 | 9 | 10 | 19 |
| 2015-F-29 | 50 | 40 | 90 |
| 2016-F11 | 5 | 2 | 7 |
| Total (*n* = 13) | 211 | 220 | 431 |
| Mean | 16.231 | 16.923 | 33.154 |
| S.D. | 18.290 | 16.276 | 33.414 |
| S.E. | 5.073 | 4.514 | 9.267 |
| Median | 8 | 10 | **17** |

Brood size more than three are shown. [a]Data presented in Narita et al. (2007b). [b]Data presented in Miyata et al. (2017). In addition to the above, four broods produced by C females were recorded; broods 2005-a, 2005-b and 2005-c contain both males and females (actual numbers not recorded) and the brood 2007-Jun-M10_f contains 37 adults, in which 7 were males and 7 were females (sex of 23 adults not recorded).
