## Supplementary material for "Persistence of a *Wolbachia*-driven sex ratio bias in an island population of *Eurema* butterflies": Table S2

Number of males and females with different *Wolbachia* infection status

| | Females | | | | Males | | | | Proportion of females | Proportion of CF females |
|---|---|---|---|---|---|---|---|---|---|---|
| | C | CF | N/A | Total | C | CF | N/A | Total | | |
| **(A) Tanegashima population** | | | | | | | | | | |
| Tanegashima (May, 2005) | 4 | 10 | 1 | 15 | 0 | 0 | 0 | 0 | 1.000 | 0.714 |
| Tanegashima (Sep, 2005) | 6 | 35 | 1 | 42 | 0 | 0 | 0 | 0 | 1.000 | 0.854 |
| Tanegashima (Oct., 2005) | 2 | 4 | 0 | 6 | 7 | 0 | 0 | 7 | 0.462 | 0.667 |
| Tanegashima (Jun, 2006) | 8 | 33 | 2 | 43 | 10 | 0 | 0 | 10 | 0.811 | 0.805 |
| Tanegashima (Sep, 2006) | 9 | 92 | 1 | 102 | 7 | 0 | 0 | 7 | 0.936 | 0.911 |
| Tanegashima (Jun, 2007) | 5 | 18 | 0 | 23 | 10 | 0 | 1 | 11 | 0.676 | 0.783 |
| Tanegashima (Sep, 2007) | 4 | 49 | 1 | 54 | 3 | 0 | 0 | 3 | 0.947 | 0.925 |
| Tanegashima (Sep, 2009) | 6 | 36 | 7 | 49 | 9 | 0 | 0 | 9 | 0.845 | 0.857 |
| Tanegashima (Jun, 2013) | 8 | 55 | 1 | 64 | 10 | 0 | 0 | 10 | 0.865 | 0.873 |
| Tanegashima (May, 2014) | 11 | 13 | 9 | 33 | 11 | 0 | 43 | 54 | 0.379 | 0.542 |
| Tanegashima (May, 2015) | 10 | 22 | 3 | 35 | 0 | 0 | 29 | 29 | 0.547 | 0.688 |
| Tanegashima (Jul, 2016) | 5 | 27 | 1 | 33 | 5 | 0 | 0 | 5 | 0.868 | 0.844 |
| Tanegashima (Aug, 2017) | 2 | 21 | 1 | 24 | 0 | 0 | 0 | 0 | 1.000 | 0.913 |
| Total | 80 | 415 | 28 | 523 | 72 | 0 | 73 | 145 | 0.865 (median) | 0.844 (median) |
| **(B) Other populations** | | | | | | | | | | |
| Zao, Miyagi Pref. (Sep, 2006) | 8 | 0 | 0 | 8 | 12 | 0 | 0 | 12 | 0.400 | |
| Tochigi, Tochigi Pref. (Sep, 2009) | 16 | 0 | 0 | 16 | 45 | 0 | 0 | 45 | 0.262 | |
| Shodoshima, Kagawa Pref. (Aug, 2011) | 32 | 0 | 0 | 32 | 0 | 0 | 103 | 103 | 0.237 | |
| Hachijojima (Sep, 2011) | 64 | 0 | 0 | 64 | 0 | 0 | 75 | 75 | 0.460 | |
| Morioka, Iwate Pref. (Aug, 2006) | 4 | 0 | 0 | 4 | 15 | 0 | 0 | 15 | 0.211 | |
| Matsudo, Chiba Pref. (Sep, 2006) | 2 | 0 | 0 | 2 | 34 | 0 | 0 | 34 | 0.056 | |
| Kawazu, Shizuoka Pref. (Sep, 2006) | 1 | 0 | 0 | 1 | 9 | 0 | 0 | 9 | 0.100 | |

N/A: not analyzed for *Wolbachia* infection.
