## Supplementary material for "Persistence of a *Wolbachia*-driven sex ratio bias in an island population of *Eurema* butterflies": Table S3

Collection record of *Eurema mandarina* on Tanegashima Island (year 1960-2007)

| Year | Northern part (Nishino-Omote) | | Middle part (Nakatane-cho) | | Southern part (Minamitane-cho) | | Total | |
|---|---|---|---|---|---|---|---|---|
| | Females | Males | Females | Males | Females | Males | Females | Males |
| 1960 | 0 | 1 | - | - | - | - | 0 | 1 |
| 1963 | - | - | 1 | 0 | - | - | 1 | 0 |
| 1969 | 0 | 1 | - | - | - | - | 0 | 1 |
| 1975 | - | - | 2 | 4 | 8 | 11 | 10 | 15 |
| 1976 | 1 | 1 | 0 | 1 | 1 | 0 | 2 | 2 |
| 1977 | 0 | 1 | 0 | 1 | 1 | 0 | 1 | 2 |
| 1978 | 1 | 3 | 1 | 0 | - | - | 2 | 3 |
| Total (1960-1978) | 2 | 7 | 4 | 6 | 10 | 11 | 16 | 24 |
| 2002 | 2 | 4 | 0 | 3 | - | - | 2 | 7 |
| 2003 | 3 | 4 | 3 | 0 | - | - | 6 | 4 |
| 2004 | 1 | 1 | 5 | 2 | - | - | 6 | 3 |
| 2005 | - | - | 0 | 1 | - | - | 0 | 1 |
| 2006 | 1 | 0 | 0 | 1 | - | - | 1 | 1 |
| 2007 | 2 | 0 | - | - | - | - | 2 | 0 |
| Total (2002-2007) | 9 | 9 | 8 | 7 | - | - | 17 | 16 |
| Grand total | 11 | 16 | 12 | 13 | 10 | 11 | 33 | 40 |

The numbers of the butterfly in the table were calculated based on the many reports of this butterfly found in Tanegashima, in the journal *SATSUMA* from Vol. 16 (1961) - Vol. 58 (2008) published by Kagoshima Entomological Society, Kagoshima, Japan, and the records of the findings by Yukiyoshi Ogata, Tanegashima, which were kindly provided to us.
