## Supplementary material for "Persistence of a *Wolbachia*-driven sex ratio bias in an island population of *Eurema* butterflies": Table S4

**Table S4**
Wing size of C females and CF females

| | Mean wing size of C females [mm] (*n* ) | Mean wing size of CF females [mm] (*n* ) | *P* |
|---|---|---|---|
| **(A) Tanegashima population** | | | |
| Tanegashima (Jun, 2006) | 23.50 ± 1.25 (*n* = 8) | 22.44 ± 1.64 (*n* = 33) | 0.09649 |
| Tanegashima (Sep, 2006) | 20.83 ± 1.27 (*n* = 9) | 20.07 ± 1.30 (*n* = 91) | 0.09383 |
| Tanegashima (Jun, 2007) | 23.90 ± 2.36 (*n* = 5) | 23.47 ± 1.53 (*n* = 18) | 0.62731 |
| Tanegashima (Sep, 2007) | 19.75 ± 1.19 (*n* = 4) | 20.34 ± 1.22 (*n* = 49) | 0.35794 |
| Tanegashima (Sep, 2009) | 20.08 ± 1.07 (*n* = 6) | 20.53 ± 1.31 (*n* = 36) | 0.43618 |
| Tanegashima (Jun, 2013) | 24.13 ± 0.92 (*n* = 8) | 23.37 ± 1.16 (*n* = 55) | 0.08565 |
| | | **Liner mixed model** | **0.04221** |
| **(B) Other populations** | | | |
| Zao, Miyagi Pref. (Sep, 2006) | 22.81 ± 1.31 (*n* = 8) | | |
| Tochigi, Tochigi Pref. (Sep, 2009) | 22.73 ± 1.43 (*n* = 15) | | |
| Shodoshima, Kagawa Pref. (Aug, 2011) | 21.81 ± 1.86 (*n* = 31) | | |
| Hachijojima (Sep, 2011) | 21.78 ± 1.32 (*n* = 62) | | |
| Morioka, Iwate Pref. (Aug, 2006) | 20.88 ± 0.85 (*n* = 4) | | |
