## Supplementary material for "Persistence of a *Wolbachia*-driven sex ratio bias in an island population of *Eurema* butterflies": Table S5

**Table S5**
Mating frequencies of C females and CF females

| Visits | No. of C females with *n* spermatophore(s) | | | | | | | Proportion of virgin in C females | Mean no. of spermatophores per C female | No. of CF females with *n* spermatophore(s) | | | | | | | Proportion of virgin in CF females | Mean no. of spermatophores per CF female |
|---|---|---|---|---|---|---|---|---|---|---|---|---|---|---|---|---|---|---|
| | *n* = 0 (virgin) | *n* = 1 | *n* = 2 | *n* = 3 | *n* = 4 | *n* = 5 | *n* = 6 | | | *n* = 0 (virgin) | *n* = 1 | *n* = 2 | *n* = 3 | *n* = 4 | *n* = 5 | *n* = 6 | | |
| Sep, 2005 | 3 | 2 | 0 | 0 | 0 | 0 | 0 | 0.600 | 0.400 | 22 | 7 | 0 | 0 | 0 | 0 | 0 | 0.759 | 0.241 |
| Jun, 2006 | 2 | 5 | 0 | 1 | 0 | 0 | 0 | 0.250 | 1.000 | 15 | 15 | 2 | 0 | 0 | 0 | 0 | 0.469 | 0.594 |
| Sep, 2006 | 2 | 7 | 0 | 0 | 0 | 0 | 0 | 0.222 | 0.778 | 30 | 41 | 18 | 0 | 0 | 0 | 0 | 0.337 | 0.865 |
| Jun, 2007 | 2 | 2 | 1 | 0 | 0 | 0 | 0 | 0.400 | 0.800 | 1 | 12 | 2 | 0 | 0 | 0 | 0 | 0.067 | 1.067 |
| Sep, 2007 | 3 | 0 | 0 | 1 | 0 | 0 | 0 | 0.750 | 0.750 | 17 | 25 | 6 | 1 | 0 | 0 | 0 | 0.347 | 0.816 |
| Sep, 2009 | 0 | 3 | 2 | 1 | 0 | 0 | 0 | 0.000 | 1.667 | 6 | 23 | 5 | 2 | 0 | 0 | 0 | 0.167 | 1.083 |
| Jun, 2013 | 4 | 3 | 1 | 0 | 0 | 0 | 0 | 0.500 | 0.625 | 31 | 21 | 1 | 0 | 0 | 1 | 0 | 0.574 | 0.519 |
| Jul, 2016 | 1 | 4 | 0 | 0 | 0 | 0 | 0 | 0.200 | 0.800 | 14 | 12 | 0 | 1 | 0 | 0 | 0 | 0.519 | 0.556 |
| Aug, 2017 | 2 | 0 | 0 | 0 | 0 | 0 | 0 | 1.000 | 0.000 | 17 | 3 | 0 | 1 | 0 | 0 | 0 | 0.810 | 0.286 |
| Total | 19 | 26 | 4 | 3 | 0 | 0 | 0 | 0.365 | 0.827 | 153 | 159 | 34 | 5 | 0 | 1 | 0 | 0.435 | 0.702 |
| | | | | | | | | 0.400 (median) | 0.778 (median) | | | | | | | | 0.469 (median) | 0.594 (median) |
